# Changes in Seasonal Footwear Elicited Alterations in Gait Kinematics but Not Stability

**DOI:** 10.1101/2023.05.18.541138

**Authors:** Sydney N. Garrah, Aaron N. Best, Amy R. Wu

## Abstract

During daily walking, humans might contend with various perturbations from slippery surfaces in the winter to uneven sidewalks in the summer. Inertial sensors enable investigations of how humans maintain balance under these natural conditions, but conducting these outdoor studies has practical considerations that might influence study results, such as the selection of footwear under different weather conditions. Our study investigates the effects of winter and summer shoe types on gait patterns, specifically whether different shoe types induce changes in gait stability measures under the same walking environment. Twelve healthy adult participants walked indoors with winter and summer shoes while their gait kinematics were recorded using an inertial sensor-based motion capture system. Spatiotemporal measures, body kinematics, stability measures (minimum margin of stability and local divergence exponent), and stepping regressions were calculated to evaluate differences between walking in summer and winter shoes. Statistical significance was determined by paired t-tests. Varying shoe types altered spatiotemporal and kinematic measures, such as increased stride time and stance time while wearing winter shoes, but increased step width and reliance on stepping were the only stability-related changes found. Our study provides insights into the influence of footwear for inertial sensor-based gait studies in real environments, aiding the analysis and interpretation of those studies to augment our understanding of natural stability behavior.

## 1. Introduction

Humans innately use multiple strategies to mitigate the risk of falling when encountering perturbations from external factors while walking. While in-lab experiments may capture elements of the ground surfaces one might experience outside [1], measuring gait under natural walking conditions might yield new insights into gait stability. Inertial sensors allow for out-of-lab motion capture studies due to their ability to accurately record motion without the need for a fixed camera system. A recent study in our lab investigated outdoor walking during winter and summer months with inertial sensors to analyze changes in the selection of stability strategies [2]. However, one confounding factor with measuring natural walking is that footwear selection would reasonably change between summer and winter conditions.

Previous studies have used inertial sensors to examine gait behavior under outdoor conditions. Schmitt et. al. investigated gait differences during indoor and outdoor walking with inertial sensors [3]. Their subjects exhibited the most variable gait while outdoors and also walked with greater walking speed, shorter double support time, and longer stride lengths outdoors compared to indoors. Another outdoor study measured gait parameters and metabolic energy expenditure from subjects walking on five different terrain types, including on dirt, gravel, and woodchips [4]. Using inertial sensors to estimate foot trajectories, they found that terrain type influenced stride variability and was correlated with energetic cost. Although inertial sensors were able to quantify gait changes over differing walking surfaces, the implied consistency in shoe type limits the variety of terrain that can be investigated. Understanding the effect of shoes on gait behavior is, therefore, of importance for elucidating gait in real world conditions. Previous walking studies on shoe types have primarily focused on gait kinematics and differences between shod and barefoot walking or shod walking with common shoe types like low or high heels [5, 6]. Few studies have examined gait stability with differing shoe types. Polome et al. investigated walking in shoes common in work environments (heeled and flat shoes) and barefoot walking to determine how footwear impacted mediolateral stability [7]. They found no significant differences in margin of stability or spatiotemporal measures (step width, step length, stance time). Menant et al. studied the effect of shoe features, including sole hardness, heel height, and tread, on stability while walking over even and uneven terrains. They found that shoe heel height and softness affected lateral balance control [8]. While these two studies did not use inertial sensors, Hollander et al. used inertial sensors to measure local dynamic stability during barefoot and shod indoor and outdoor walking [6]. They found no changes in stability for shoe types. All these studies were either performed indoors (overground or on a treadmill) or outdoors on flat ground. Hence, their conclusions on gait stability do not reflect the breadth of outdoor terrain one might experience and the accompanying practical shoe requirements.

Here, we investigated the effects of winter and summer shoe types on gait patterns using inertial sensors in a controlled indoor setting. Specifically, our study aims to clarify which gait changes, if any, can be attributed to shoe changes, instead of terrain surface conditions. We anticipated that summer shoes could be represented by trainers or sneakers while boots with traction would be typical winter shoes. Since winter boots are not expected to have high heels or soft soles, we hypothesized that comparing winter and summer footwear would find no differences in spatiotemporal and stability measures and thus no differences in stability strategies.

## 2. Materials and Methods

To measure differences between walking in summer and winter types of shoes, healthy adult subjects (*N* =12, five males, seven females, weight70.2±9.6 kg, height 171.1±6.0 cm, age from 21 to 24 years) performed two indoor walking trials – one with their own winter shoes and one with summer shoes (Figure 1). For each trial, subjects walked overground for five minutes at their preferred speeds on a predetermined rectangular path, averaging approximately 135±13 strides (mean±s.d.) strides per trial after the removal of turns. Gait kinematics were recorded using an inertial sensor-based motion capture system (MVN Link, Xsens, Netherlands) with seventeen inertial sensors on all segments of the body (Figure 1). We recorded the last three minutes of each trial to allow two minutes for acclimatization to each shoe type. The order of shoe type was randomized for each subject. Full details on data processing and analysis can be found in our previous paper [2], and we provide a short summary of our methods here.

**Figure 1:**
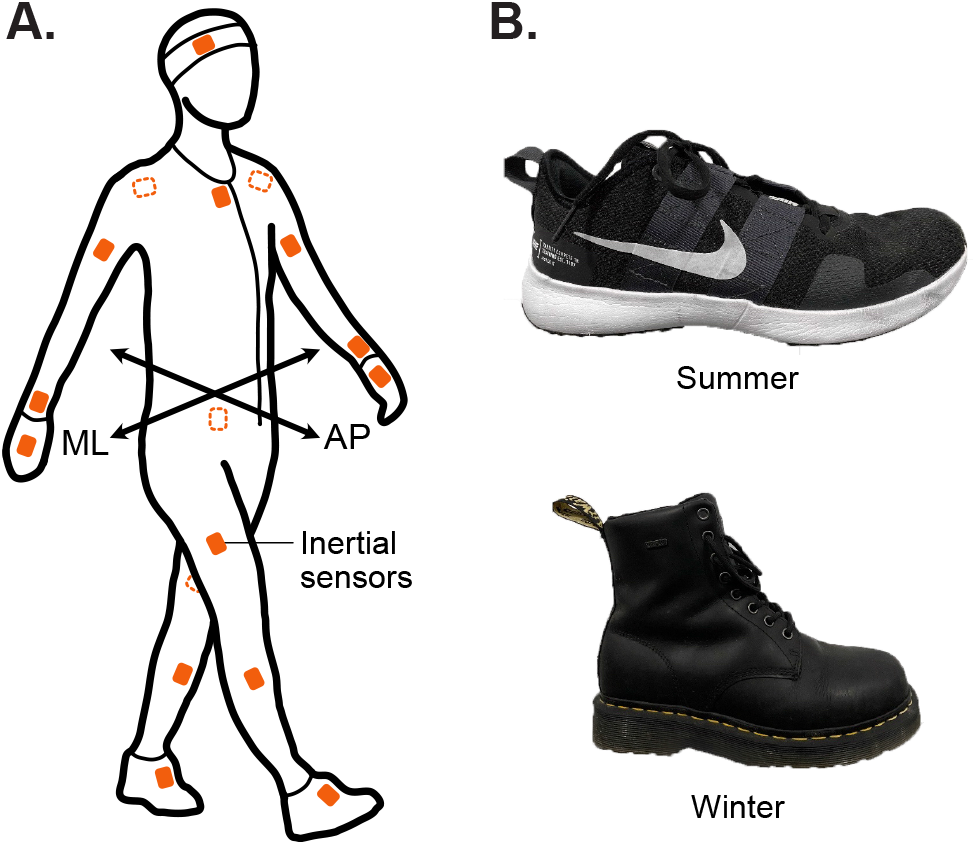
Experimental setup and representative shoes worn by subjects. Kinematic data was captured from (A) an inertial sensor suit worn by subjects as they walked in either (B) summer shoes (top) or winter shoes (bottom). ML: mediolateral; AP: anterior-posterior

Spatiotemporal measures, body kinematics, and three stability measures (margin of stability, local divergence exponent, stepping regression) were calculated to compare walking in the two different shoe types. Spatiotemporal measures included walking speed, stride time, stance time, stride length, and step width and their root-mean-square (RMS) variability. We also measured body center of mass (CoM) kinematics and trunk angles to observe differences in body kinematics. We compared the range of the CoM position and velocity in the anterior-posterior (AP), mediolateral (ML), and vertical (VT) directions. Walking speed was determined from the average AP CoM velocity. We also evaluated the mean and range trunk flexion (AP) and lateral trunk lean. A trunk angle of zero denotes that the trunk is perpendicular to the ground.

For stability measures, we chose the minimum Margin of Stability (MoS), the Local Divergence Exponent (LDE), and stepping regression goodness-of-fit (*R*^2^). The MoS aims to quantify the relationship between the CoM and the foot contact with the ground. Minimum MoS was defined as the minimum distance between the extrapolated CoM and the base of support (BoS) [9]. In the ML direction, we defined the outer bound of the BoS as the lateral component of the fifth metatarsal position on the stance foot and calculated the minimum MoS during single support. In the AP direction, we used the heel position to define the BoS because the minimum difference between the BoS and XCoM occurs at heel strike for the AP direction [10]. If the AP and ML XCoM positions exceed the BoS, the MoS is considered negative. The fifth metatarsal and heel positions were estimated by Xsens.

The LDE is a measure of whole body stability by computing the average logarithmic rate of divergence of a system after a small perturbation [11]. It can be calculated using any source of kinematic data, and we elected to use CoM velocity in the AP, ML and VT direction. The creation of a state space and calculation of the LDE was completed using an open-source version of the algorithm proposed by Mehdizadeh [12, 13]. A larger LDE indicates a more chaotic and less stable system. We used the short-term exponent *λ*_*s*_ as our LDE measure.

Stepping regressions measure the natural step-to-step variation during steady walking [14]. Inferring that the position of the foot during a step is dependent on the previous kinematic behaviour of the pelvis, regressions have been used to evaluate the reliance of foot placement on the state of the upper body. We performed a linear regression between the deviation of the state of the CoM at midstance and the deviation of the location of the subsequent heel strike in both the ML and AP direction. The goodness-of-fit (*R*^2^) value was compared between shoe types to determine if the relationship between pelvis kinematics and foot placement varied between shoe conditions.

For each measure, paired t-tests were used to determine statistical significance (*α* = 0.05). If a result was significant, its effect size (*η*^2^) was calculated [15]. Prior to statistical comparisons, all measures were normalized by standing leg length *L* (defined as the distance from the ground to the greater trochanter of the femur) and gravitational acceleration *g* as base units. Stride length was normalized by *L* (mean 0.908 m), stride time by 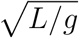 (mean 0.304 s), speed by 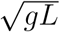 (mean 2.98 ms^*−*1^), and acceleration by *g*. The study was approved by the university Institutional Review Board, and subjects provided informed consent prior to the experiment.

## 3. Results

We found some differences between winter and summer shoes, primarily in the spatiotemporal measures and body kinematics compared to stability-related measures (summarized in Table 1). Spatiotemporal and kinematic changes with winter shoes included longer stride time, wider step width, larger range of ML CoM position and velocity, and greater forward trunk lean with winter shoes (Figure 2 and 3). In contrast, no significant differences were found for MoS and LDE stability measures (Figure 4A and B), but there was a greater reliance on ML stepping behaviour with winter shoes (Figure 4C).

**Table 1:** Spatiotemporal, kinematic, and stability measures (mean±s.d.) for walking with winter shoes and summer shoes. *P* -values indicate statistical significance between shoe types (\**P <* 0.05) with statistically significant results accompanied by their respective effect size (*η*^2^). Trunk angles are reported in degrees. All other quantities are reported in dimensionless form with leg length and gravity as base units. RMS: root-mean-square, MoS: margin of stability, LDE: Local Divergence Exponent

| Measure | | Winter | Summer | $P$ ($\eta^2$) |
| --- | --- | --- | --- | --- |
| Walking Speed | Mean | $0.44 \pm 0.04$ | $0.45 \pm 0.05$ | 0.201 |
| Stride Time | Mean | $3.65 \pm 0.19$ | $3.53 \pm 0.19$ | 0.002* (0.105) |
| Stance Time | Mean | $2.15 \pm 0.13$ | $2.08 \pm 0.13$ | 0.002* (0.077) |
| Stride Length | Mean | $1.60 \pm 0.09$ | $1.58 \pm 0.11$ | 0.175 |
| Step Width | Mean | $0.05 \pm 0.04$ | $0.03 \pm 0.03$ | 0.045* (0.070) |
| Walking Speed | RMS | $0.02 \pm 0.006$ | $0.01 \pm 0.002$ | 0.135 |
| Stride Time | RMS | $0.07 \pm 0.02$ | $0.06 \pm 0.01$ | 0.140 |
| Stance Time | RMS | $0.05 \pm 0.02$ | $0.04 \pm 0.008$ | 0.096 |
| Stride Length | RMS | $0.04 \pm 0.02$ | $0.03 \pm 0.004$ | 0.159 |
| Step Width | RMS | $0.054 \pm 0.007$ | $0.049 \pm 0.008$ | 0.010* (0.106) |
| AP CoM Position | Range | $1.59 \pm 0.09$ | $1.57 \pm 0.11$ | 0.187 |
| ML CoM Position | Range | $0.050 \pm 0.008$ | $0.046 \pm 0.008$ | 0.003* (0.076) |
| VT CoM Position | Range | $0.044 \pm 0.006$ | $0.043 \pm 0.006$ | 0.128 |
| AP CoM Velocity | Range | $0.10 \pm 0.01$ | $0.09 \pm 0.01$ | 0.006* (0.074) |
| ML CoM Velocity | Range | $0.12 \pm 0.02$ | $0.11 \pm 0.02$ | 0.008* (0.043) |
| VT CoM Velocity | Range | $0.14 \pm 0.02$ | $0.15 \pm 0.02$ | 0.821 |
| AP Trunk Angle | Mean | $4.71 \pm 1.71$ | $3.75 \pm 1.93$ | 0.019* (0.069) |
| ML Trunk Angle | Mean | $-0.34 \pm 0.74$ | $-0.23 \pm 0.61$ | 0.488 |
| AP Trunk Angle | Range | $3.32 \pm 0.71$ | $3.20 \pm 0.58$ | 0.191 |
| ML Trunk Angle | Range | $3.30 \pm 1.09$ | $3.36 \pm 0.87$ | 0.734 |
| AP MoS | Mean | $-0.12 \pm 0.04$ | $-0.13 \pm 0.05$ | 0.299 |
| ML MoS | Mean | $0.02 \pm 0.02$ | $0.01 \pm 0.02$ | 0.178 |
| AP LDE | Mean | $1.84 \pm 0.18$ | $1.77 \pm 0.12$ | 0.096 |
| ML LDE | Mean | $1.71 \pm 0.114$ | $1.69 \pm 0.07$ | 0.624 |
| VT LDE | Mean | $1.45 \pm 0.12$ | $1.47 \pm 0.10$ | 0.404 |
| AP Regression $R^2$ | Mean | $0.90 \pm 0.05$ | $0.88 \pm 0.04$ | 0.274 |
| ML Regression $R^2$ | Mean | $0.84 \pm 0.04$ | $0.81 \pm 0.04$ | 0.011* (0.101) |

**Figure 2:**
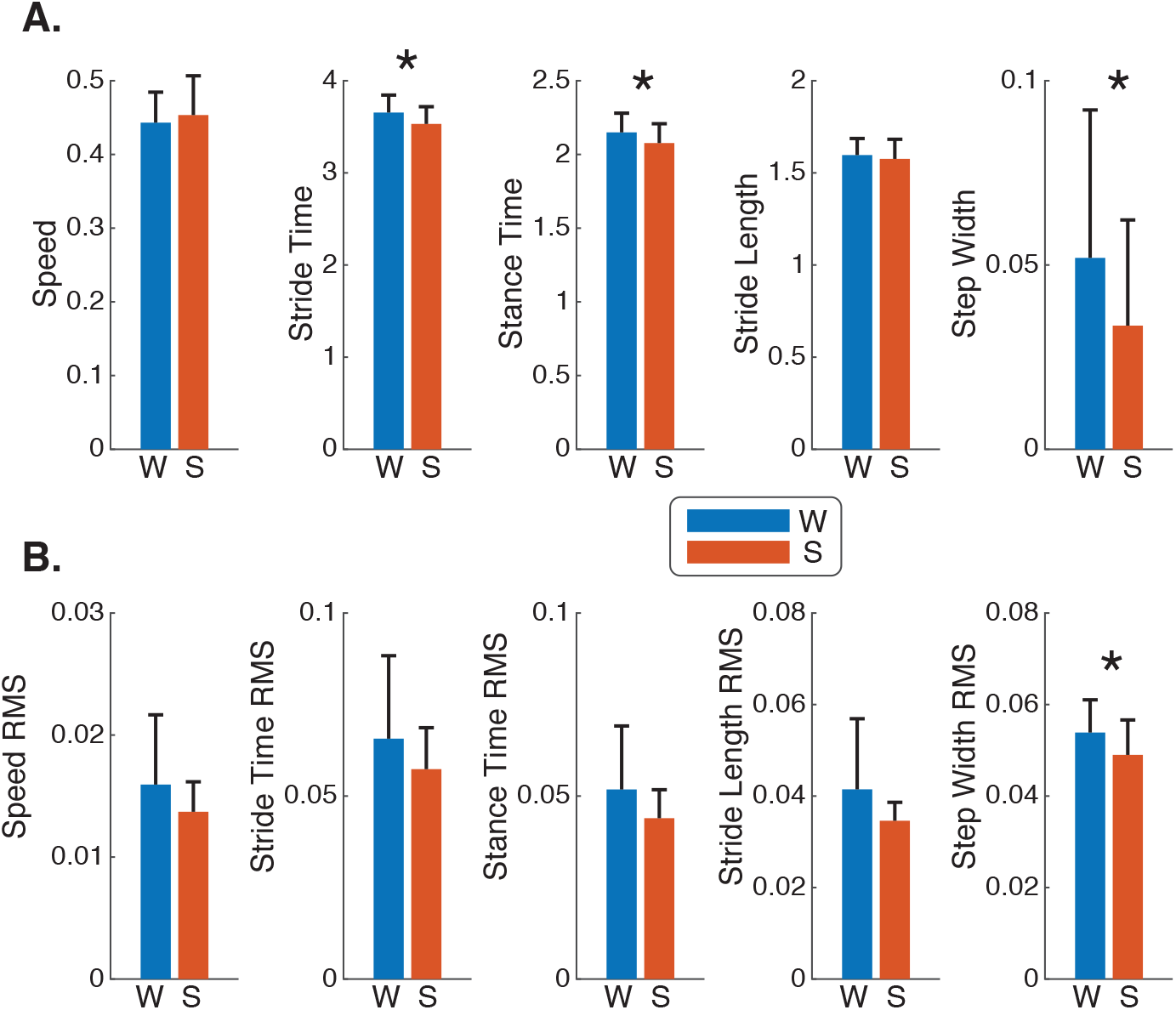
Spatiotemporal measures for walking in winter and summer shoes. (A) Mean and (B) root-mean-square (RMS) variability measures for walking speed, stride time, stance time, stride length, and step width. Bars denote averages across all subjects (*N* = 12), and error bars denote one standard deviation. Statistical significance between shoe types indicated by an asterisk (\**P <* 0.05).

**Figure 3:**
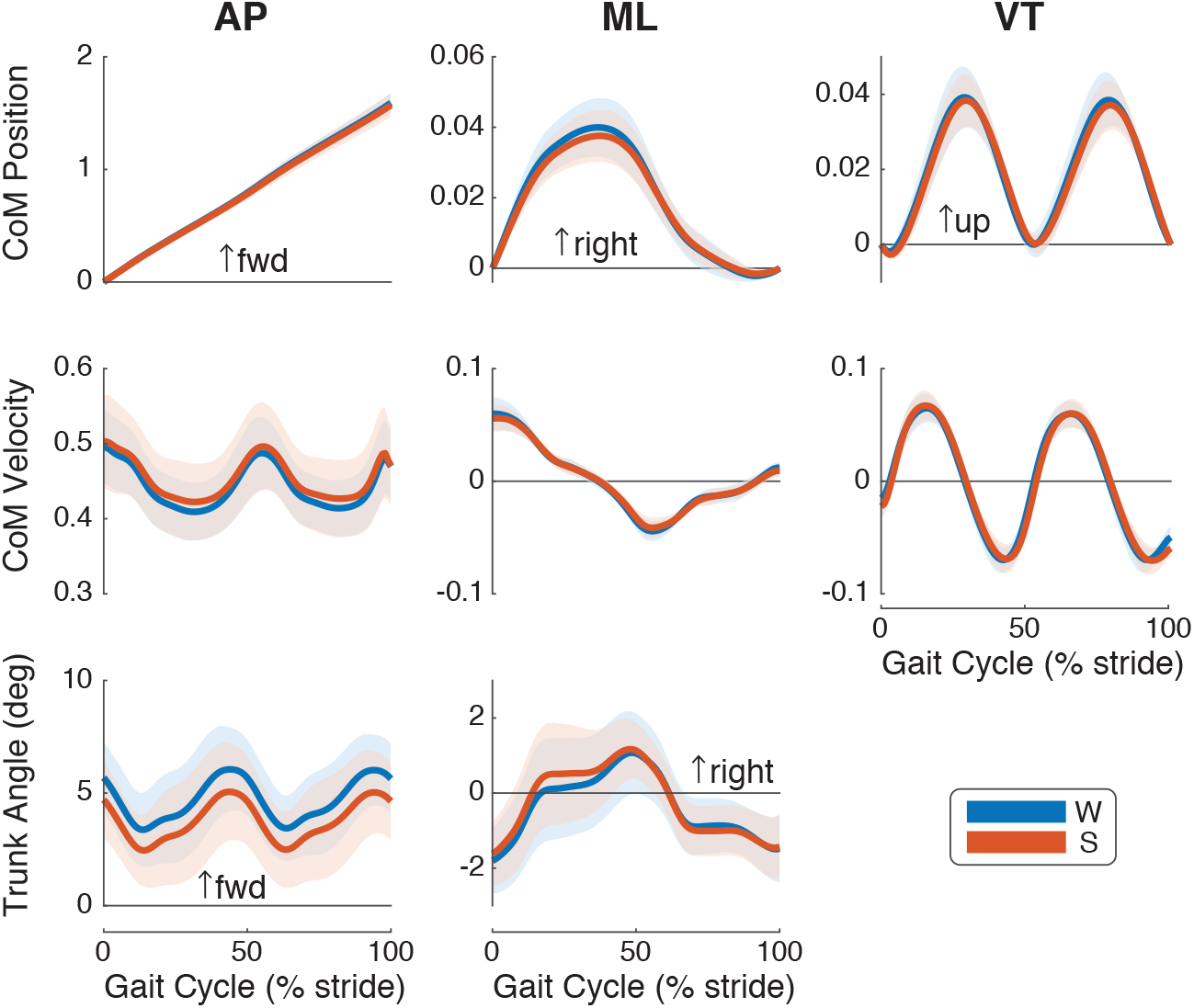
Body CoM kinematics and trunk angle over a stride. Mean CoM position, CoM velocity, and trunk angles of all subjects (*N* = 12) with standard deviation indicated by shaded region. The gait cycle (horizontal axis) is shown as a percentage of stride starting with heelstrike. CoM kinematics are shown in dimensionless form.

**Figure 4:**
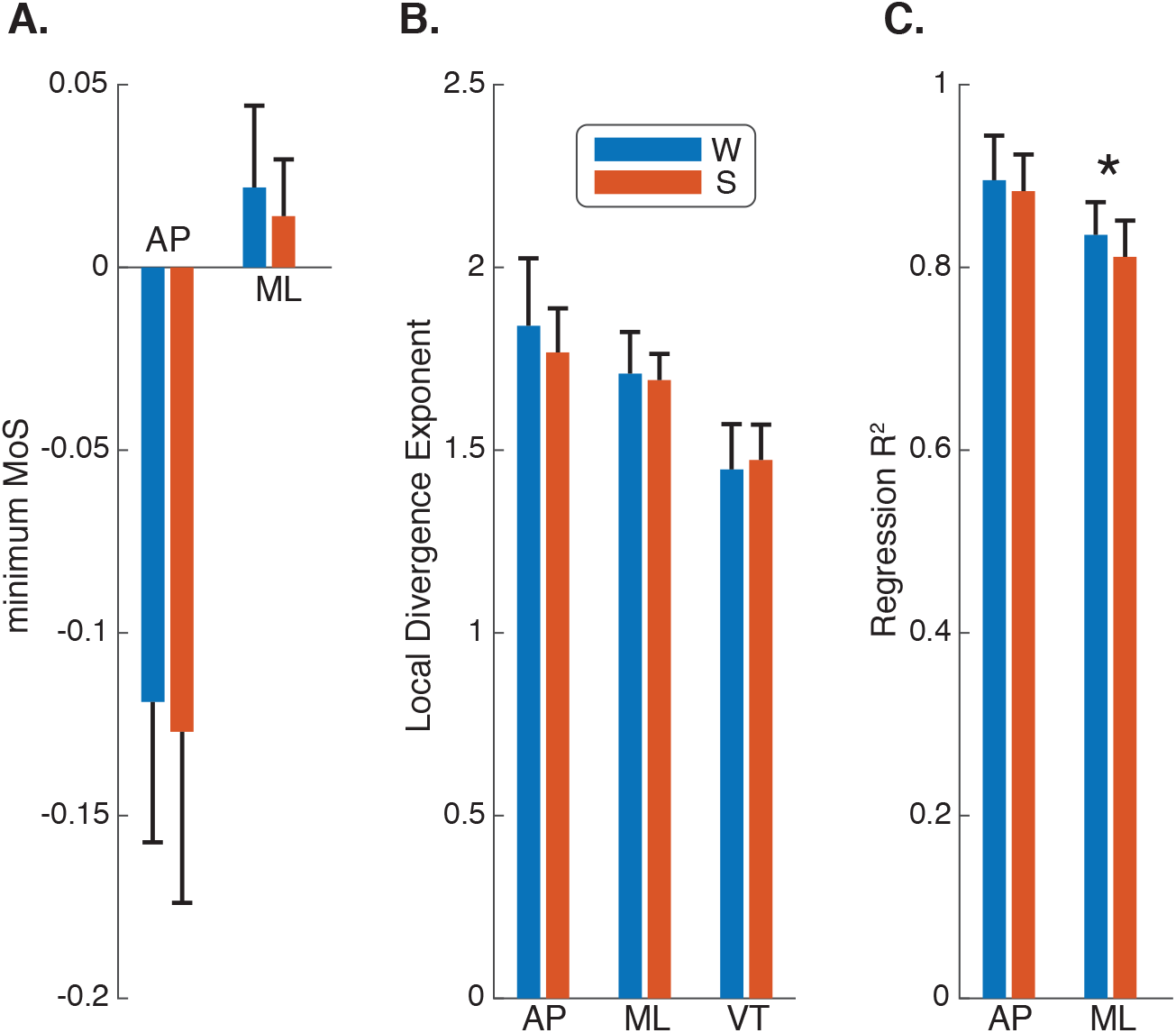
Gait stability while walking in winter and summer shoes, as measured by (A) the minimum MoS (AP, ML), (B) Local Divergence Exponent, or LDE, (AP, ML, VT), and (C) regression goodness-of-fit *R*^2^ (AP, ML). Bars denote averages across all subjects (*N* = 12), and error bars denote one standard deviation. Statistical significance between shoe types indicated by an asterisk (\**P <* 0.05).

Subjects exhibited spatiotemporal changes between the two walking conditions. We found that walking in winter shoes led to approximately 3.5% longer stride (*P* = 0.002, *η*^2^ = 0.105) and stance times (*P* = 0.002, *η*^2^ = 0.077) and 55% greater step width (*P* = 0.045, *η*^2^ = 0.070) compared to summer shoes (Figure 2). Winter shoes also led to a 10% increase in step width variability (*P* = 0.010, *η*^2^ = 0.106). No changes were found in walking speed, stride length, or other tested stride variability measures (Table 1).

We also found differences in kinematic measures between shoe types (Figure 3, Table 1). The range of motion for ML CoM position and velocity increased with winter shoes by 9.1% (*P* = 0.003, *η*^2^ = 0.076) and 7.4% (*P* = 0.008, *η*^2^ = 0.043), respectively. AP CoM velocity range also increased by 7.9% with winter shoes (*P* = 0.006, *η*^2^ = 0.074). Subjects also exhibited 0.95^*◦*^ more trunk flexion with winter shoes than summer shoes (*P* = 0.019, *η*^2^ = 0.069).

Shoe types did not affect subject stability, as measured by MoS and LDE, but did alter their reliance on the stepping strategy (Figure 4, Table 1). We found no significant differences for minimum MoS between shoe conditions in the AP (*p* = 0.299) or ML (*p* = 0.178) direction. Similarly, the LDE stability measure was also not significantly different in all three planes of motion between shoe types (*p* = 0.096, *p* = 0.624, and *p* = 0.404 for the AP, ML and VT directions, respectively). While the *R*^2^ value for the AP stepping regression was also not significantly different (*p* = 0.274), the *R*^2^ value for ML regression was approximately 3% greater with winter shoes than summer shoes (*p* = 0.011, *η*^2^ = 0.101).

## 4. Discussion

Our study sought to elucidate the effects of typical summer and winter shoes on gait using inertial sensors. We hypothesized that no spatiotemporal, stability, or stepping behaviour differences would be found between these two footwear types. Our stability results agreed with our hypothesis with no significant changes found for MoS and LDE. However, we did find an increased reliance on the stepping strategy in the ML direction. Spatiotemporal measures, such as stride time and step width, were also influenced by shoe type as well as ML CoM position and velocity range.

We found that footwear had a greater effect on spatiotemporal parameters and body kinematics than on stability measures. Footwear seemed to more strongly influence gait in the ML direction with increased step width, step width variability, CoM position and velocity ranges, and reliance on the stepping strategy with winter shoes.

Although we did not expect differences between tested shoe types, winter shoes did alter spatiotemporal and kinematic behaviour compared to summer shoes. Winter shoes led to longer stride times, which seemed influenced by greater stance time. Thus, subjects had longer foot contact with the ground while walking with winter shoes. Perhaps winter shoes were heavier than summer shoes, leading to a reduction in swing phase. Alternatively, the additional traction on winter shoes could have increased the friction between the shoe and the ground, also resulting in longer stance times. Winter shoes also led to more lateral CoM sway (i.e., range of ML CoM position), accompanied by a larger velocity range. The lateral sway could be related to the wider step widths by both stepping wider and moving the body more laterally with each step. Subjects walked at similar speeds (mean 1.34 ms^*−*1^) with both shoe types, but AP CoM velocity also exhibited a larger range. However, it is unclear if the 8% increase between summer (mean range 0.33 ms^*−*1^) and winter (mean range 0.35 ms^*−*1^) is meaningful or not. Subjects also exhibited more forward trunk lean, which may be related to exerting additional effort to walk in possibly heavier winter shoes.

Despite the increase in the ML CoM range of position and velocity in the winter shoes condition, the ML MoS was not significantly different due to the simultaneous increase in the step width. We also found no significant difference in the AP MoS, and no significant difference in the LDE in any direction. Although the measures of stability remained similar between the different conditions, subjects exhibited behaviour that indicated changes in, or a greater need for, stabilization strategies. Wider step widths and greater step width variability indicated increased control of ML stepping behaviour. These stepping results are aligned with the ML stepping regression *R*^2^ results. These three changes (larger step widths, step width variability, and ML *R*^2^) indicate some potential defense against instability which did not manifest with MoS and LDE measures.

Although previous studies had differing types of footwear, our study both contradicted and supported their findings. Our results contradict the findings of Polome et al. who found no effect of footwear on stance time [7]. A study from de Jong et al. investigated spatiotemporal and stability measures between orthopedic and non-orthopedic shoes in subjects with Hereditary Motor Sensory Neuropathy [16]. Although this study focused on a different subject group than the current study, no differences in stability between in the two shoe types were found. However, the study found improved gait with orthopedic footwear, defined as increased gait speed and step length and decreased step time and step width [16]. Our results, along with those of the de Jong et al., suggest that spatiotemporal measures can be altered by footwear without necessarily eliciting a change in stability.

Our study has several limitations. We found no changes with the MoS and LDE stability measures, but it is possible that other stability measures might yield significant changes. Although wider step widths was the only significant stability-related change, inertial sensors cannot directly measure distance between each other, and estimates of step width may be less accurate than other spatiotemporal estimates [17, 18, 19]. In addition, the number of strides available for the LDE analysis was 109 strides, and more strides could lead to more precise estimates of *λ*_*s*_ [20]. Although we found several significant changes in different gait parameters, the accompanying effect sizes were primarily medium-sized (∼0.07) [21] with the largest effect sizes (∼0.10) for stride time, step width RMS, and ML regression *R*^2^. Walking speed remained unchanged, which might not be reflective of traversing over slippery winter terrain. Because we aimed to quantify the effect of practical footwear changes when conducting outdoor gait studies, we did not enforce that subjects wear the same shoes. However, self-selected shoes for the summer and winter were similar among subjects (e.g. running shoes for summer, boots for winter, see Data Availability for shoe images) with the exception of one subject who wore sandals as summer shoes. For both shoe types, we placed an inertial sensor midline at the top of the foot (instep). It is possible that sensor re-placement after changing shoe type could affect our results. However, any differences are expected to be minor due to instep placement compared to other possible sensor locations [22].

Inertial sensors provide a unique opportunity to investigate gait in diverse environments, leading to greater scientific understanding of how humans maintain balance while walking. However, practical considerations, such as proper footwear in icy or snowy conditions, might affect study conditions. Our study provides insights into the influence of footwear for inertial sensor-based gait studies in real environments. Our findings suggest that gait stability is not significantly altered by changes in footwear and is more likely to be controlled by other external factors, such as the walking environment. Therefore, stability changes elicited between walking during winter and summer months are more likely a consequence of ground conditions rather than shoe type. As more and more gait studies take place outside, studies such as ours can help to differentiate between uncontrolled factors of interest (e.g., walking surface) and those of less or no interest (e.g., shoe type or clothing), leading to deeper insights into how people naturally behave and maintain stability in real life.

## Author Contributions

ANB and ARW conceived the study, and SNG and ANB designed the study. SNG acquired and analyzed the data. SNG and ARW drafted the article. ANB and ARW provided supervision and critical feedback. ARW provided funding. All authors provided data interpretation, critically revised the article, and approved of the final version.

## Data Availability

Study data can be accessed at Queen’s University Dataverse repository at [The Effect of Summer and Winter Footwear on Gait Kinematics and Stability](https://borealisdata.ca/privateurl.xhtml?token=30bc4459-b6f0-42f0-8d6b-493f2b9f1c19).

## Acknowledgements

Funding was provided by the Natural Sciences and Engineering Research Council of Canada (NSERC) Discovery Grant (grant no. RGPIN-2019-05221), NSERC Undergraduate Summer Research Award, NSERC Alexander Graham Bell Canada Graduate Scholarships - Doctoral Program (CGS-D), and Ingenuity Labs Research Institute.

